# Hormonal and Environmental Drivers of Spermiation in the Endangered Mountain Yellow-Legged Frog (*Rana muscosa*): Toward Biologically Informed Assisted Reproductive Technologies

**DOI:** 10.1101/2025.08.21.671667

**Authors:** Natalie E. Calatayud, Leah Jacobs, Rose Upton, Stephanie Chancellor, Barbara S. Durrant, Debra M. Shier

## Abstract

Amphibians are among the most threatened vertebrates, yet assisted reproductive technologies (ARTs) remain underutilised in their conservation. We developed and evaluated a biologically relevant, non-lethal sperm collection protocol for *Rana muscosa*, a critically endangered frog in a long-term conservation breeding program. Thirteen hormone treatments were tested across six post-injection time points, and sperm quality was assessed via concentration, motility, osmolality, and pH. Generalised linear mixed models revealed that gonadotropin-releasing hormone (GnRHa) alone significantly outperformed hCG-based regimens. The 3 µg/g GnRHa dose yielded the highest sperm concentration and sustained motility from 3 to 24 hours post-injection. Motility was highest under moderately acidic (pH 6.5–7.0) and hypoosmotic (75–100 mOsm/kg) conditions. To support decision-making, we applied a refined Wildlife Sperm Index (WSI) that weights motility and concentration as primary functional-yield traits, with motility given greater relative weight given its direct role in fertilisation, and treats pH and osmolality as secondary environmental-compatibility traits. Under this revised composite score, 3 µg/g GnRHa achieved the highest overall WSI score, outperforming 2 µg/g and 4 µg/g GnRHa, which ranked second and third, respectively, despite their more favourable pH and osmolality profiles. These findings provide the first evidence-based ART protocol for *R. muscosa* and offer a transferable framework for optimising gamete collection, IVF, and cryopreservation in line with other amphibian species, advancing both genetic management and species recovery goals.

## 1. Introduction

Global conservation frameworks increasingly recognise that preserving genetic diversity is critical to halting biodiversity loss. The IUCN’s One Plan Approach exemplified this integration by promoting both in situ and ex situ strategies for species recovery (https://www.cpsg.org/our-work/our-approach/one-plan-approach). Building on this foundation, the current Amphibian Conservation Action Plan (IUCN SSC Amphibian Specialist Group, 2024) emphasises the importance of cryopreservation, assisted reproductive technologies (ARTs), and genome resource banking (GRB) as vital tools to strengthen and extend One Plan implementation, particularly for amphibians. The Convention on Biological Diversity’s Global Biodiversity Framework (Target 4) calls for the maintenance of genetic diversity through both *in situ* and *ex situ* measures, which, while not explicitly stated, can reasonably be interpreted to encompass the development and use of genetic repositories (Convention on Biological Diversity, 2022). These priorities are particularly urgent for amphibians, the most threatened vertebrate class for which nearly half of all known species are in decline, and over 40% are threatened with extinction (IUCN SSC Amphibian Specialist Group, 2024; Luedtke et al., 2023; Stuart et al., 2004). Yet, amphibians remain underrepresented in ex situ conservation efforts. Only ∼3% of threatened amphibian species are maintained in zoological institutions through conservation breeding programs (CBPs), and even fewer are represented in genome resource banks (hereafter referred to as biobanks) (Conde et al., 2013, 2011; IUCN SSC Amphibian Specialist Group, 2024). However, these strategies (CBPs and biobanks) are now widely recognised as essential for amphibian species, not only for safeguarding against extinction but also for bolstering fragmented or declining wild populations, especially in the face of emerging infectious diseases, such as chytridiomycosis (Clulow et al., 2019; IUCN SSC Amphibian Specialist Group, 2024).

Assisted reproductive technologies (ARTs), including hormone-induced gamete induction, artificial fertilisation, and cryopreservation, offer critical support for conservation breeding. When paired with CBPs, ARTs can enhance reproductive output, reduce the need for wild collection of individuals, and improve genetic management (Catenazzi, 2015; Clulow et al., 2019; Conde et al., 2013, 2011; Convention on Biological Diversity, 2022; Della Togna et al., 2020; Howell et al., 2021a, 2021b; IUCN SSC Amphibian Specialist Group, 2024; Luedtke et al., 2023; Stuart et al., 2004). The updated ACAP recognises ARTs as vital to amphibian recovery and encourages species-specific protocols to improve breeding efficiency, genetic retention, and long-term storage (IUCN SSC Amphibian Specialist Group, 2024). Although ARTs cannot replace in situ conservation efforts, they offer a powerful complement, particularly in small, declining populations, where inbreeding, reduced fertility, and ageing founders threaten long-term viability. In cases where wild population numbers are too low to sustain recovery in situ, ARTs and other ex situ tools become essential for preventing extinction and facilitating future reintroductions. Non-lethal gamete retrieval and cryopreservation can help maintain reproductive potential while minimising the impact on remnant wild populations (Clulow and Clulow, 2016).

The mountain yellow-legged frog (*Rana muscosa*) exemplifies the urgency and complexity of developing ARTs for conservation. Listed as endangered by both the California Department of Fish and Wildlife and the United States Fish and Wildlife Service, *R. muscosa* now persists in only a few isolated populations across three mountain ranges in southern California. Since 2006, the San Diego Zoo Wildlife Alliance (SDZWA) has managed a CBP for *R. muscosa*, releasing more than ∼15,000 individuals into the wild. Despite this progress, challenges persist: founder genetic diversity remains extremely limited (Steiner et al., 2024), breeding success is inconsistent (Calatayud et al., 2022; Jacobs et al., 2021), and reliance on a narrow subset of breeding pairs may further erode genetic health over time.

To improve reproductive success and enable long-term genetic management, we sought to develop a biologically relevant, non-lethal sperm collection protocol for *R. muscosa*. Because amphibians with external fertilisation are highly sensitive to their surrounding medium, we also evaluated environmental factors affecting sperm quality. Osmotic pressure and extracellular pH are particularly influential: sperm motility is typically activated under hypoosmotic conditions (<100 mOsm/L), peaks between 0–50 mOsmol/L, and declines as osmolality increases (Browne et al., 2015; Byrne et al., 2015; Otero et al., 2023; Reinhardt et al., 2015; Silla et al., 2017). Similarly, sperm activation is suppressed under acidic conditions and optimised near neutral-to-slightly alkaline pH, with even larger shifts (0.2–0.5 units) affecting dynein-ATPase activity, ion transport, and flagellar function (Hamamah and Gatti, 1998; Nishigaki et al., 2014; Zhou et al., 2015). For *R. muscosa*, maintaining optimal osmolality and pH during gamete collection is likely essential to maximise sperm viability and fertilisation success.

This study presents a systematic evaluation of hormone-induced spermiation in *Rana muscosa*, with six primary objectives: (1) develop a biologically relevant, non-lethal protocol for sperm collection via hormone induction; (2) evaluate the efficacy of different doses and combinations of gonadotropin-releasing hormone (GnRHaa; D-Ala, des-Gly¹ ethylamide) and human chorionic gonadotropin (hCG); (3) quantify sperm output based on concentration and motility; (4) identify optimal post-injection time windows for sperm collection; (5) examine how environmental covariates (pH and osmolality) affect sperm activation, motility, and viability; and (6) apply a Wildlife Sperm Index (WSI) to rank hormone treatments based on combined biological and environmental parameters (Jacobs et al., 2025). This approach enables a comprehensive assessment of sample quality, ensuring that selection is informed by the combined performance across relevant biological and environmental parameters rather than isolated measures.

## 2. Materials and Methods

### 2.1 Standard Housing and Care

An ex-situ population of *R. muscosa* was maintained at the SDZWA’s Beckman Center for Conservation Research as part of a long-term conservation breeding program. Frogs were housed in temperature-controlled, recirculating aquatic systems equipped with mechanical and biological filtration, water chillers, and UV sterilisation. Each enclosure was outfitted with basking platforms, polyresin caves, and artificial foliage for environmental enrichment.

To simulate in situ seasonal cycles, water temperatures were reduced to 3–4 °C during winter brumation (December–March) and gradually increased to 19 °C in summer. Photoperiods were adjusted weekly using programmable timers to match natural daylength patterns from the San Jacinto Mountains. Water quality was monitored 2–3 times weekly and adjusted as needed using reconstituted reverse osmosis water. Frogs were fed gut-loaded crickets and small larvae 3–5 times per week during active periods and were fasted during brumation and the breeding season.

### 2.2 Hormone Protocols and Sperm Collection

To evaluate hormone-induced spermiation, we tested 13 hormone treatments and a saline control (Simplified Amphibian Ringer; SAR) (Table 1). Hormone regimens included different doses of gonadotropin-releasing hormone agonist (GnRHa; BACHEM H-4070) and human chorionic gonadotropin (hCG; Sigma CG5-10VL), administered either alone or in combination (Table 1). Hormones were delivered via intraperitoneal injection in the early morning. A baseline urine sample (0 h) was collected immediately prior to injection, and post-treatment samples were collected at 1, 3, 5, 7, and 24 hours.

**Table 1.** Number of male frogs, total injections, and sample counts across repeated time points for each treatment over the 10-year study period. Number of unique male frogs, total injections, and total sample counts across time points (0, 1, 3, 5, 7, and 24 h) for each hormone treatment. Males (n = 170; aged 3–12 years) were randomly assigned to one of 13 hormone treatments or a vehicle control using a randomised block design to control for age and individual variability. Over the 10-year study period, 437 injections were administered, with 63 males reused across multiple trials following a minimum 4-week inter-treatment interval to prevent carryover effects. Each injection event involved repeated sampling at multiple time points, so sample counts represent events rather than unique individuals.

| Dosage | Number of males | Total injections | Sample count across time points (0, 1, 3, 5, 7 24 h) |
| --- | --- | --- | --- |
| SAR (control) | 21 | 53 | 305 |
| hCG 5 IU | 10 | 24 | 324 |
| hCG 10 IU | 15 | 32 | 144 |
| hCG 5 IU / g + GnRHa 0.3 µg / g | 11 | 25 | 286 |
| hCG 5 IU / g + GnRHa 0.6 µg / g | 9 | 22 | 198 |
| hCG 10 IU / g + GnRHa 0.3 µg / g | 11 | 28 | 360 |
| hCG 10 IU / g + GnRHa 0.6 µg / g | 9 | 20 | 110 |
| 0.3 µg / g GnRHa | 12 | 28 | 161 |
| 0.6 µg / g GnRHa | 14 | 29 | 123 |
| 1µg/g GnRHa | 10 | 18 | 198 |
| 2 µg/g GnRHa | 14 | 36 | 306 |
| 3 µg/g GnRHa | 12 | 34 | 234 |
| 4 µg/g GnRHa | 12 | 30 | 162 |
| **TOTAL** | **160** | **437** | **2,911** |

*2.6 Statistical Analysis*

Sperm samples were collected non-lethally and assessed for concentration and motility. A minimum of 10 spermatozoa per sample was required for motility analysis (see below for the rationale for the criteria). Sperm induction trials were conducted in a seasonally concentrated manner: spring-summer (March–June) and fall (September–November) to capture potential seasonal variation. No trials were conducted during winter brumation (December–February).

### 2.3 Experimental Design and Animal Assignment

We used a randomised block design to assign 170 unique male frogs (aged 3–12 years) to hormone treatments, controlling for individual variability and age. Over the course of the study (2015-2017 & 2019-2024), 437 hormone injections were administered. Sixty-three frogs were reused across multiple trials, with a minimum 4-week interval between injections to prevent carryover effects. Each injection event involved repeated sampling at multiple time points (Table 1). Because individuals could contribute to multiple trials, the total sample size reflects events rather than unique individuals.

### 2.4 Sample Collection Procedures

Urine was collected using the cradle-and-girdle technique, in which frogs were held gently by the forelimbs over a Petri dish to stimulate urination. If urination did not occur, a sterile vinyl catheter (0.34 mm x 0.052 mm, Scientific Commodities #BB31785-V/5) was inserted into the cloaca to retrieve fluid. Samples were transferred to microcentrifuge tubes for analysis and refrigerated until evaluation.

### 2.5 Sperm Quality Assessment

Each urine sample was evaluated for: (1) sperm concentration, (2) percent motility, (3) osmolality (Osm), and (4) pH (Table S1). Sperm concentration was assessed using a Bright Line haemocytometer (Millipore-Sigma #Z359629) and phase contrast microscope (Nikon Eclipse 80i). For each sample, 10 μL of urine was loaded and manually counted. Viscous samples were diluted 1:5 or 1:10 in distilled water, and the final concentration was calculated by adjusting for dilution. Sperm motility was assessed using established methods (Calatayud et al., 2025; Langhorne et al., 2021). Motility was calculated as the percentage of motile sperm (mot/[mot + non-motile]) based on samples with ≥ 10 sperm (See *Sperm Motility by Hormone and Time* below). Osmolality and pH were measured using a vapour pressure osmometer and calibrated pH meter, respectively.

### 2.6 Statistical Analysis

All analyses were conducted in R version 4.2.1 using RStudio. The dataset comprised repeated measures collected from individual male *Rana muscosa* subjected to exogenous hormone treatments. Two sperm quality metrics were analysed: sperm concentration and sperm motility, measured as raw motile and non-motile sperm counts. Overdispersion was accounted for by including frog ID (id) as a random effect in all models. Analytical stages are detailed below.

#### 2.6.1 Sperm Concentration by Hormone and Time

Sperm concentration was modelled using a Tweedie-distributed generalised linear mixed model (GLMM) with a log link, implemented via the glmmTMB package (Brooks et al., 2017). The Tweedie family was selected to accommodate non-negative, right-skewed data with overdispersion and a high frequency of zeros. Hormone treatment (trt) and time post-injection (time) were treated as categorical fixed effects, and their interaction (trt × time) was explicitly included to test for time-specific treatment effects. Only biologically relevant time points—0, 1, 3, 5, 7, and 24 hours post-injection—were retained. Frog ID (id) was included as a random effect.

Model assumptions and goodness of fit were assessed using the DHARMa package (Hartig, 2022) to evaluate the residual distribution, dispersion, and potential zero inflation. Estimated marginal means (EMMs) and 95% confidence intervals were extracted using the emmeans package (Lenth, 2025) and back-transformed from the log scale. Hormone treatments were ranked by overall model-estimated sperm concentration. To identify time-specific peaks in sperm concentration, an iterative peak-finding algorithm adapted from Della Togna et al. (2017) was applied (Della Togna et al., 2017). At each step, time points exceeding the mean plus one standard deviation (SD) of the remaining time points were classified as peaks, removed from the dataset, and the process was repeated until no further peaks were identified.

#### 2.6.2 Sperm Motility by Hormone and Time

Sperm motility was modelled using a zero-inflated binomial generalised linear mixed model (ZIB-GLMM) with a logit link, implemented via the glmmTMB package (Brooks et al., 2017). The response variable was specified as a two-column matrix of raw counts, cbind(mot, nm), where mot and nm denote motile and non-motile sperm, respectively. This model structure allowed for distinction between biological zeros (e.g., presence of sperm but no motility) and structural zeros (e.g., no sperm available for assessment). Hormone treatment (trt) and time post-injection (time) were treated as categorical fixed effects, and their interaction (trt × time) was included to evaluate temporal variation in treatment response. Individual frog ID (id) was incorporated as a random effect to account for repeated measures. A zero-inflation component (ziformula = ∼1) was included to explicitly model excess zeros associated with biologically meaningful motility failure.

Only observations in which ≥10 sperm were observed were included in the motility analysis. These cut-off balances statistical robustness with biological relevance. From a methodological perspective, using fewer than 10 observations yields unstable binomial proportion estimates with wide confidence intervals. At a sample size of 10, the maximum standard error for estimated motility is approximately 15.8%, which was deemed acceptable for comparative modelling. Biologically, this threshold is equivalent to observing 10 sperm in 0.1 µL — the volume encompassed by the 25 central squares of a standard Neubauer haemocytometer. This extrapolates to ∼1 million sperm per mL, a widely accepted benchmark for functional fertility assessment in amphibians.

Model fit and assumptions were evaluated using the DHARMa package (Hartig, 2022). Estimated marginal means (EMMs) and 95% confidence intervals were calculated on the response scale using the emmeans package (Lenth, 2025) and back-transformed to predicted percent motility. Pairwise comparisons were conducted to estimate odds ratios (ORs) for hormone treatments across time points. For treatment-level contrasts, odds ratios were computed across all pairwise combinations of hormone regimens, collapsed across time. For time-specific analyses, odds ratios were calculated for pairwise comparisons of motility at each time point within treatments, and for interactions between specific treatment × time combinations. Data wrangling and visualisation were performed using the dplyr and ggplot2 packages (Wickham, 2016).

#### 2.6.3 Environmental Influences – pH and Osmolality

To assess whether pH and osmolality (Osm) were influenced by hormone treatment, these variables were analysed as response variables using Gaussian generalised linear mixed models (GLMMs). Hormone treatment (trt) was treated as a categorical fixed effect, and frog ID (id) was included as a random effect. Models were implemented using the glmmTMB package (Brooks et al., 2017). Because we were restricted by sample volumes, only samples with complete pH and Osm values were retained. Model residuals were visually inspected to confirm normality and homoscedasticity. Estimated marginal means (EMMs) and pairwise contrasts used to calculate odds ratios (ORs) were compared between hormone treatments using the emmeans package (Lenth, 2025). Visualisations were generated using ggplot2 (Wickham, 2016).

To evaluate the influence of environmental parameters on sperm concentration, we modelled the effects of pH and osmolality (Osm) using both generalised linear and additive approaches. Analyses were limited to samples from animals treated with 2, 3, or 4 µg/g GnRHa, and included only observations with complete pH, Osm, and concentration values. Sperm concentration was modelled using a Tweedie GLMM with a log link, with pH and Osm included as fixed effects and frog ID as a random effect, implemented via glmmTMB (Brooks et al., 2017). To assess potential nonlinearities and interactions, we also fit a generalised additive model (GAM) using the mgcv package (Wood, 2017). In the GAM, pH was modelled as a linear term, Osm as a smoothed term (s(Osm_scaled)), and their interaction was included as a tensor product (ti(pH_scaled, Osm_scaled)), with a random intercept for individual identity (s(id, bs = “re”)). Prior to modelling, pH and Osm values were centred and scaled using dplyr (Wickham 2023). Predictions were generated across a biologically relevant grid of pH (6.5–8.0) and Osm (0–100 mOsm/kg), back-transformed to the response scale, and expressed as model-estimated sperm concentration. Modelling treated both pH and Osm as continuous variables, whereas heatmaps and estimated marginal means (EMMs) were based on biologically informed bins to aid interpretation.

To model the effect of environmental conditions on motility, both generalised linear and generalised additive models were used. Analyses were restricted to animals treated with 2, 3, or 4 µg/g GnRHa. Observations were filtered to include only those with complete data for motility, pH, and Osm, and with a total sperm count ≥10. A zero-inflated binomial GLMM was fit using glmmTMB (Brooks et al., 2017), with pH and Osm as continuous fixed effects and frog ID as a random effect. EMMs were estimated for combinations of pH (6.5, 7.0, 7.5) and Osm (200, 250, 300 mOsm/kg), selected based on observed ranges and prior amphibian literature, using emmeans (Lenth, 2025). Odds ratios were calculated for all pairwise contrasts. To explore nonlinear responses and interactive effects, a GAM was fit using mgcv (Wood, 2017). pH was modeled as a linear term; Osm was modeled as a smooth term (s(Osm scaled)); and their interaction was specified as a tensor product (ti(pH_scaled, Osm_scaled)). Frog ID was modelled as a random intercept (s (id, bs = “re”)). Prior to modelling, pH and Osm values were centred and scaled (dplyr). Model predictions were generated across a grid of pH (6.5–8.0) and Osm (0–100 mOsm/kg), then back-transformed to the response scale and expressed as percent motility.

#### 2.6.4 Wildlife Sperm Index (WSI)

To rank overall sample quality across treatments, we utilized the Wildlife Sperm Index (WSI) (Jacobs et al., 2025) [23] to reflect the functional relationship between sperm traits and fertilisation potential. Rather than treating concentration, motility, pH, and osmolality as four independently assessed components, we distinguished functional-yield traits (concentration and motility), which directly determine fertilisation potential, from environmental-compatibility traits (pH and osmolality), which influence sperm function without themselves constituting fertility outcomes. The WSI does not currently incorporate acrosome integrity, DNA integrity, or mitochondrial function, as these require additional processing and specialised assays beyond the scope of this field-based, non-lethal protocol; concentration, motility, pH, and osmolality were retained as core inputs because they can be assessed rapidly and non-destructively from the same urine sample.

Within the functional-yield term, concentration was weighted more only slightly more heavily than motility, reflecting the importance of obtaining sufficient amounts of sperm while incorporating motility’s more direct role in fertilisation. Within the environmental-compatibility term, pH was weighted more heavily than osmolality, given its substantially larger main effect on motility (Estimate = 1.14, p < 0.001) versus osmolality’s (Estimate = 0.162, p < 0.001). All traits were scaled on a 0-100 scale prior to scoring. Because per-sample WSI scores were strongly right-skewed, treatment differences were assessed using a Kruskal-Wallis test followed by pairwise Mann-Whitney U tests with Bonferroni correction.

## 3. RESULTS

### 3.1 Sperm Concentration by Hormone and Time

Sperm concentration varied significantly across hormone treatments and time points (Tweedie GLMM: treatment, χ² = 248.6, df = 12, p < 0.001; time, χ² = 120.5, df = 5, p < 0.001, Table 2). Among all treatments, 3 µg/g GnRHa consistently elicited the strongest response (see Table 2 for raw means), peaking at 7 hours post-injection (hpi) and maintaining high concentrations from 3 to 24 hpi (Figure 1G). This treatment significantly outperformed all others over the 24-hour period (OR range: 2.48–3.71, p < 0.001), with model-estimated concentrations exceeding those of SAR and all hCG-containing regimens (Tables 2 and 3).

**Figure 1.** Mean sperm concentration and percentage motility over time across 12 hormone treatments. Each panel (A–L) displays bar plots of mean sperm concentration (cells/mL) at 0, 1, 3-, 5-, 7-, and 24-hours post-injection, overlaid with a dashed line and points representing mean percentage motility. Bars are colored using a log10-scaled viridis colour gradient corresponding to sperm concentration. Error bars indicate 95% confidence intervals. Motility was plotted only when sperm were present, and total sperm assessed ≥10. Treatments are as follows: A. 5 IU hCG; B. 10 IU hCG; C. 0.3 µg/g GnRHa; D. 0.6 µg/g GnRHa; E. 1 µg/g GnRHa; F. 2 µg/g GnRHa; G. 3 µg/g GnRHa; H. 4 µg/g GnRHa; I. 5 IU hCG + 0.3 µg/g GnRHa; J. 5 IU hCG + 0.6 µg/g GnRHa; K. 10 IU hCG + 0.3 µg/g GnRHa; L. 10 IU hCG + 0.6 µg/g GnRHa.

**Table 2.** Ranked summary of sperm concentration by hormone treatment of the top four treatments. Includes raw and model-estimated means (sperm/mL), standard errors), and 95% confidence intervals (CI).

| Rank | Hormone Treatment (GnRHa) | Raw Mean (sperm/mL) | SE (Raw) | 95% CI (Raw) | Model-Estimated Mean (sperm/mL) | SE (Model) | 95% CI (Model) |
| --- | --- | --- | --- | --- | --- | --- | --- |
| 1 | 3 µg/g | $1.70 \times 10^6$ | $2.38 \times 10^4$ | $1.23 \times 10^6 - 2.17 \times 10^6$ | $7.32 \times 10^5$ | $1.02 \times 10^4$ | $5.59 \times 10^5 - 9.59 \times 10^5$ |
| 2 | 2 µg/g | $5.27 \times 10^5$ | $1.21 \times 10^4$ | $2.29 \times 10^5 - 7.65 \times 10^5$ | $1.32 \times 10^5$ | $2.20 \times 10^4$ | $9.52 \times 10^4 - 1.83 \times 10^5$ |
| 3 | 4 µg/g | $4.80 \times 10^5$ | $9.60 \times 10^4$ | $2.92 \times 10^5 - 6.68 \times 10^5$ | $6.62 \times 10^4$ | $1.28 \times 10^4$ | $4.56 \times 10^4 - 9.64 \times 10^4$ |
| 4 | 1 µg/g | $1.63 \times 10^5$ | $3.81 \times 10^4$ | $8.85 \times 10^4 - 2.38 \times 10^5$ | $2.82 \times 10^4$ | $9.5 \times 10^3$ | $1.57 \times 10^4 - 5.09 \times 10^4$ |

**Table 3.** Spermiation peaks in the top three hormone treatments. Peaks were identified using the SD-threshold method, where a time point was classified as a peak if its mean concentration exceeded the mean of remaining values +1 standard deviation, iteratively.

| Hormone Treatment | Peak Time Points (hpi) | Peak Sperm Concentration (cells/mL) |
| --- | --- | --- |
| 2 µg/g GnRHa | 1, 7 | 4.39e+0<sup>5</sup>, 1.72e+0<sup>6</sup> |
| 3 µg/g GnRHa | 7 | 2.56e+0<sup>6</sup> |
| 4 µg/g GnRHa | None | None |
| 1 µg/g GnRHa | None | None |

The 2 µg/g GnRHa treatment showed a biphasic response, with prominent peaks at 1 and 7 hpi and significantly elevated concentrations compared to baseline and non-peak time points (OR = 3.12–5.91, p < 0.01). This pattern indicates a repeatable window of sperm release with multiple collection opportunities. In contrast, the 4 µg/g GnRHa treatment produced a single moderate peak at 3 hpi, with concentrations declining sharply thereafter. The likelihood of achieving higher concentrations was significantly greater at 3 hpi than at later time points (OR = 2.5–6.7, p < 0.05), indicating a narrow and early collection window. The 1 µg/g GnRHa dose produced moderate sperm output, while lower doses (0.3 and 0.6 µg/g GnRHa) yielded sparse responses with high non-responder frequency and were therefore excluded from further analysis.

All hCG-only and hCG+GnRHa combination treatments elicited poor responses across time points, with model-estimated concentrations significantly lower than those of the 2–4 µg/g GnRHa group (OR range = 0.03–0.15, p < 0.001). No sperm release was observed in SAR (control).

### 3.2 Sperm motility by hormone and time

Sperm motility varied significantly across hormone treatments and time points (Zero-Inflated Binomial GLMM: treatment, χ² = 88.0, df = 12, p < 0.001; time, χ² = 34.8, df = 5, p < 0.001; treatment × time interaction, χ² = 196.81, df = 12, p < 2.2e–16). Among all treatments, the 3 µg/g GnRHa dose consistently elicited the highest and most sustained motility, with model-estimated values exceeding 90% between 3–7 hours post-injection (hpi) and peaking at 94.5% (SE = 1.59%; 95% CI: 91.0–96.9%) at 5 hpi (Figure 1G; Table 3–5). This treatment significantly outperformed most hCG-containing regimens across the time series (OR range = 2.14–23.8, p < 0.001), with the strongest contrast observed between 3 µg/g GnRHa and 5 IU hCG + 0.3 µg/g GnRHa (OR = 23.8, p < 0.001).

**Table 4.** Summary of model-estimated mean sperm motility across all time points (1–24 hpi) for the top-performing hormone treatments. Values represent the average predicted percentage motility, standard deviation (SD), and standard error (SE) derived from zero-inflated binomial GLMM outputs.

| Hormone Treatment | Mean % Motility | SD | SE | # Time Points Measured | Peak Time Points (hpi) | Peak % Motility |
| --- | --- | --- | --- | --- | --- | --- |
| 3 $\mu\text{g/g}$ GnRHα | 63.03 | 19.40 | 8.67 | 5 | none | none |
| 4 $\mu\text{g/g}$ GnRHα | 61.37 | 15.44 | 6.91 | 5 | 1, 3, 5 | 67.96%,<br>61.43%,<br>64.52% |
| 2 $\mu\text{g/g}$ GnRHα | 61.86 | 11.88 | 5.31 | 5 | none | none |
| 1 $\mu\text{g/g}$ GnRHα | 57.76 | 5.90 | 2.63 | 5 | 1 | 64.54% |

**Table 5.** Model-estimated pH and Osmolality values by hormone treatment. SAR was used as the model intercept and reference level for comparisons. Δ values reflect model-estimated differences from SAR for each treatment.

| Hormone Treatment | Estimated pH ± SE | ΔpH vs SAR | p-value (pH) | Estimated Osm ± SE (mOsm/kg) | ΔOsm vs SAR | p-value (Osm) |
| --- | --- | --- | --- | --- | --- | --- |
| 1 μg/g GnRH<sub>a</sub> | 7.516 ± 0.094 | +0.031 | 0.763 | 52.811 ± 4.977 | +3.66 | 0.438 |
| 2 μg/g GnRH<sub>a</sub> | 7.555 ± 0.094 | +0.070 | 0.467 | 52.560 ± 4.977 | +3.40 | 0.446 |
| 3 μg/g GnRH<sub>a</sub> | 7.597 ± 0.094 | +0.112 | 0.517 | 63.173 ± 4.977 | +14.02 | 0.072 |
| 4 μg/g GnRH<sub>a</sub> | 7.424 ± 0.094 | -0.061 | 0.530 | 51.392 ± 4.977 | +2.24 | 0.623 |

The 2 µg/g GnRHa treatment followed closely, peaking at 96.6% motility at 5 hpi and maintaining high values throughout the 3–7 hpi window (Figure 1F). Compared to hCG-only treatments, the 2 µg/g dose yielded significantly higher odds of motility (OR = 3.5–7.6, p < 0.001). The 1 µg/g GnRHa treatment reached its maximum response earlier (95.5% at 3 hpi; Figure 1H) but declined more rapidly thereafter. Across the full-time series, it remained significantly more effective than combination treatments (OR range = 2.4–5.2, p < 0.01). The 4 µg/g dose displayed greater temporal variability, with motility estimates ranging between 59–76% (Figure 1E), and lower odds of success relative to the 3 µg/g treatment (OR = 0.45–0.50, p < 0.001).

Odds ratios for individual time points reinforced the dominance of GnRHa-only regimens. At 1 hpi, 3 µg/g GnRHa resulted in significantly greater odds of motility than all other treatments (OR = 2.1–2.5, p < 0.05). Similar contrasts were observed at 3 and 5 hpi, highlighting consistent temporal advantages for this treatment (Table S5).

In contrast, all hCG-only and hCG+GnRHa combination regimens elicited poor motility responses across time. Many individual samples were excluded due to insufficient sperm for scoring (<10 sperm/sample). Although 0.3 and 0.6 µg/g GnRHa elicited high motility at isolated time points, small sample sizes (1–3 observations per time point) precluded their inclusion in final analyses. The 10 IU hCG treatment was excluded due to repeated failure to meet biological and statistical thresholds for modelling. Raw mean motility model estimates are reported (Table 4 and Table S6).

Notably, motility rankings did not align with sperm concentration outcomes. While 3 µg/g GnRHa produced the highest sperm concentrations (Figure 1G), its overall motility across time was moderate (61.4%; Table 4; Table S6). By contrast, treatments such as 0.3 µg/g GnRHa and 5 IU hCG showed high motility (≥67%) but performed poorly in terms of sperm concentration. Of the four top motility regimens, only 2 µg/g GnRHa consistently ranked statistically high in both sperm motility and concentration (Figure 1; Tables 3 and 4).

### 3.3 Environmental Drivers of Sperm Output – pH and Osmolality

Hormone treatments did not significantly affect sample pH or osmolality (GLMMs; all p > 0.37), allowing for evaluation of their independent influence on sperm traits (see Table 5). Model-estimated pH values were consistent across groups (mean range: 7.42–7.60), and while osmolality varied moderately (e.g., 49.2 ± 18.2 mOsm/kg in SAR vs. 63.2 ± 18.4 mOsm/kg in the 3 µg/g GnRH group), no pairwise differences were statistically significant after Tukey correction. A Tweedie GLMM revealed a modest negative correlation between osmolality and sperm concentration (Estimate = –0.020, p = 0.029), and a model including a pH × osmolality interaction term showed a significant negative interaction (Estimate = –0.044, p = 0.018), indicating reduced sperm output at high values of both variables. A complementary GAM confirmed this pattern, with a linear effect of pH, a smooth effect of osmolality, and a significant interaction (ti(pH, Osm), p < 0.001), indicating that sperm concentration was highest at low pH and low osmolality, with diminished output at higher values of either parameter.

Sperm motility was best explained by a zero-inflated binomial GLMM, which showed significant positive main effects of pH (Estimate = 1.14, p < 0.001) and osmolality (Estimate = 0.162, p < 0.001), with a strong negative interaction between the two (pH:Osm Estimate = –0.021, p < 0.001). A generalised additive model (GAM) including the same variables yielded nearly identical results, with strong statistical support for the main effects and interaction term (ti(pH, Osm), p < 0.001). However, empirical observations did not align with the broad high-performance region predicted by the model. Instead, observed motility was sharply concentrated in a single bin—pH 7.0 and osmolality 50–100 mOsm/kg—where motility reached 73%. All other bins showed markedly reduced values (<40%, many <1%), with no evidence of gradual improvement across the modelled gradient. This suggests that motility may respond to environmental thresholds rather than continuous gradients, and that model extrapolations may overestimate motility under conditions not strongly represented in the data. This discrepancy is illustrated in Figure 2, which compares observed and predicted percent motility across the pH–osmolality landscape.

**Figure 2.** Predicted sperm motility across pH and osmolality gradients based on a generalised additive model (GAM) with a linear × smooth interaction. Observed (left) and predicted (right) sperm motility across pH and osmolality gradients for *Rana muscosa*. Observed values represent mean percent motility within discrete pH and osmolality bins. Predicted values were generated from a generalised additive model (GAM) with a linear × smooth tensor interaction (pH × s(Osm)), using scaled inputs and converted to percent motility. The model predicts highest motility (yellow) at low pH (6.5–7.0) and moderate to high osmolality (75–100 mOsm/kg). However, observed values show a more restricted pattern, with peak motility (73%) only at pH 7.0 and an osmolality of 50–100 mOsm/kg, and lower motility under all other conditions. Both panels use a viridis colour palette to facilitate direct comparison.

### 3.4 Treatment Ranking and Integration of Sperm Assessments and Micro-environmental Drivers

When incorporating environmental parameters using the revised WSI, which weights motility and concentration as primary functional-yield traits and pH and osmolality as secondary environmental-compatibility traits, 3 µg/g GnRHa achieved the highest mean WSI score (Table 6), followed by 2 and 4 µg/g GnRHa. A Kruskal-Wallis test confirmed a significant overall difference in WSI scores among the four GnRHa doses (χ² = 101.42, df = 3, p < 2.2 × 10 ^16^). Pairwise Mann-Whitney U tests with Bonferroni correction indicated that 1 µg/g GnRHa scored significantly lower than 2 µg/g (p = 3.0 × 10 ^14^), 3 µg/g (p < 2 × 10 ^16^), and 4 µg/g GnRHa (p = 5.7 × 10 ^10^), but no significant differences were detected among 2, 3, and 4 µg/g GnRHa themselves (2 vs. 3: p = 1.000; 2 vs. 4: p = 1.000; 3 vs. 4: p = 0.055). This indicates that while 3 µg/g GnRHa produced the highest descriptive WSI score, its advantage over 2 and 4 µg/g GnRHa was not statistically significant, and all three doses represent comparably effective protocols under the revised composite scoring framework.

**Table 6.** Performance summary of the top three hormone treatments (2, 3, and 4 µg/g GnRHa) based on combined sperm concentration, motility, pH and Osmolality. Estimated mean sperm concentration and percent motility were derived from model-based predictions. Values were rescaled to the range [0, 100] to standardise across metrics. A composite performance index (WSI) was calculated as a weighted combination of functional-yield traits (concentration and motility, combined multiplicatively) and environmental-compatibility traits (pH and osmolality). Treatments are ranked by descending combined score. The 3 µg/g GnRHa treatment achieved the highest average WSI score, reflecting its combined advantage in sperm concentration and motility, despite a less favourable pH and osmolality profile than 4 µg/g GnRHa.

| Hormone Treatment (GnRHa µg/g) | Mean Concentration (cells/mL) | Mean % Motility | Range Osmolality | Range pH | Average WSI |
| --- | --- | --- | --- | --- | --- |
| 3 | 732,105 | 63.03 | 19 -150 | 7.0-8.1 | 21.19 |
| 2 | 66,249 | 61.86 | 11 - 122 | 6.5-8.5 | 18.79 |
| 4 | 132,029 | 61.37 | 2 - 147 | 6.4-8.5 | 15.56 |

## 4. Discussion

This study presents a comprehensive evaluation of hormone-induced spermiation in *Rana muscosa*, a critically endangered amphibian reliant on conservation breeding. By integrating hormonal, temporal, and environmental parameters, we developed a biologically relevant, non-lethal sperm collection protocol to support assisted reproductive technologies (ARTs), including cryopreservation, in vitro fertilisation (IVF), and long-term genetic management.

Among the tested protocols, GnRHa-only treatments significantly outperformed hCG-only and combination regimens, consistent with findings in many other externally fertilising amphibians (Clulow et al., 2019; Otero et al., 2023; Silla et al., 2021). The 3 µg/g GnRHa dose produced the highest sperm concentration and sustained motility, making it the most effective overall. The 2 and 4 µg/g doses also induced strong responses and offer practical alternatives depending on collection timing or individual variability. In contrast, hCG, which has been found to be optimal for some species (Clulow et al., 2018; Silla and Byrne, 2019; Upton et al., 2024), failed to produce consistent or statistically usable sperm samples, highlighting species-specific differences in hormonal sensitivity.

Sperm concentration and motility both peaked between 5–7 hours post-injection, establishing this as the optimal collection window in the better hormone protocols. The broader temporal range observed for 3 µg/g GnRHa (3–24 hpi) offers logistical flexibility for facility or field-based applications. Relative to other externally fertilising amphibians (e.g., *Crinia*, *Litoria*, *Pseudophryne* spp.), captive *R. muscosa* have lower mean sperm concentrations and more variable motility following hormonal induction (Browne et al., 2015; Silla et al., 2017). These differences may reflect species-specific reproductive physiology, but also likely result from long-term captivity, inbreeding, or underlying endocrine dysfunction, highlighting the need for reproductive monitoring of species before significant inbreeding, where possible. Notably, testicular abnormalities and cases of pseudo-hermaphroditism have been observed in this species (Jacobs et al., 2021), which could contribute to inconsistent hormone responsiveness and compromised sperm quality.

Environmental conditions strongly influenced sperm motility. Optimal motility occurred under moderately acidic to neutral pH conditions (pH 6.5–7.0) and hypoosmotic media (75–100 mOsm/kg) relative to the typical osmolality of amphibian plasma (230–250 mOsm/kg), aligning with known activation thresholds in other externally fertilising amphibians (Byrne et al., 2015; Silla and Byrne, 2019). Although hormone treatments did not significantly alter pH or osmolality, variation across samples highlights the importance of environmental control during sperm collection and fertilisation procedures.

Our analyses identified a significant negative interaction between pH and osmolality on sperm motility, indicating that optimal activation cannot be achieved by adjusting either factor in isolation. At the cellular level, extracellular pH modulates several pathways, including dynein ATPase activity, intracellular calcium signalling, and ion transport mechanisms (e.g., Na/H exchangers, bicarbonate transporters), that regulate sperm flagellar function and motility initiation (Hamamah and Gatti, 1998; Nishigaki et al., 2014; Zhou et al., 2015). Osmolality further interacts with these pathways by controlling hydration and ionic balance during activation. Broadly across taxa—including mammals, fish, amphibians, and even insects—sperm function is highly sensitive to deviations in environmental osmolality and pH, with activation and longevity often restricted to narrow, species-specific ranges (Alavi and Cosson, 2006; Browne et al., 2019; Byrne et al., 2015; Reinhardt et al., 2015). In externally fertilising vertebrates, especially anurans, sperm are often activated at extremely low osmolalities and show longer motility durations compared to fish, likely reflecting evolutionary adaptation to freshwater spawning environments. However, optimal performance varies not only between species but among populations, shaped by both physiological constraints and local environmental pressures. These findings underscore the need for precisely defined, taxon-specific sperm handling protocols when developing assisted reproductive technologies, as even modest mismatches in pH or osmolality can profoundly impact fertilisation potential.

To guide decision-making across multiple biological endpoints, we applied a revised Wildlife Sperm Index (WSI), distinguishing functional-yield traits (concentration and motility) from environmental-compatibility traits (pH and osmolality), and weighting motility most heavily given its direct role in fertilisation. This framework allowed us to rank treatments according to both sperm output and environmental compatibility, while ensuring that high concentration alone could not compensate for poor motility. Under this revised structure, 3 µg/g GnRHa achieved the highest descriptive WSI score, combining the highest raw sperm concentration with motility comparable to the other GnRHa doses, despite a less favourable pH and osmolality profile than 4 µg/g GnRHa; however, this advantage was not statistically significant relative to 2 or 4 µg/g GnRHa, indicating that all three doses perform comparably once concentration and motility are appropriately prioritised as the primary determinants of fertilisation potential. This illustrates the importance of how composite metrics like the WSI are constructed: the relative weighting of functional versus environmental traits can materially affect which protocol is identified as the top performer, underscoring the need for biologically grounded, rather than arbitrary, weighting schemes when applying multi-trait indices to reproductive management decisions. An important next step will be to validate the WSI against actual reproductive outcomes, such as fertilisation success following IVF or post-thaw sperm function in cryopreserved samples, to confirm whether higher WSI scores reliably predict greater fertility potential and more suitable material for biobanking. Without this validation, the WSI remains a useful tool for ranking relative sample quality, but its capacity to forecast actual reproductive success has not yet been empirically tested.

## 5. Conclusions

This study provides direct applications for the recovery of R. muscosa. Fertility challenges within the conservation breeding program, including inconsistent breeding, poor responses to environmental cues, and suspected reproductive pathologies, have constrained program sustainability (Jacobs et al., 2021; Steiner et al., 2024). Using a revised Wildlife Sperm Index that prioritises functional sperm traits over environmental compatibility, 3 µg/g GnRHa was identified as the most broadly effective hormone protocol, combining the highest sperm concentration with motility comparable to other GnRHa doses, although this advantage was not statistically distinguishable from 2 or 4 µg/g GnRHa. The protocol developed here enables non-lethal sperm collection from genetically valuable males, targeted sperm retrieval for IVF or cryopreservation, and species-specific benchmarks for hormone dosing and activation conditions. Several biological and logistical variables remain unresolved. Sperm output and quality may vary with individual health, age, duration in captivity, or genetic diversity, factors not explicitly tested here. Future research should also evaluate additional sperm quality characteristics, such as DNA degradation, acrosome integrity, and mitochondrial function, validate the WSI against actual fertilisation and biobanking outcomes, and examine the effects of genetic diversity and other environmental stressors on sperm quality. Strategic crosses may further improve genetic diversity and reproductive output. More broadly, the approach developed here is applicable to other externally fertilising amphibians, particularly those in conservation breeding programs facing seasonal breeding constraints, small founder sizes, and low reproductive success. Standardised, evidence-based ART protocols, like the one presented here, are critical for scaling up cryobanking and IVF efforts in amphibian recovery initiatives.

## Acknowledgements

All research was permitted by the U.S. Fish and Wildlife Service recovery permit 76006B-3, an MOU from the California Department of Fish and Wildlife and approved by the Institutional Animal Care and Use Committee (IACUC protocols #15-00, 18-003), in accordance with institutional, national, and international guidelines. This study also adhered to the U.S. Fish and Wildlife Service scientific permit TE-76006B-0. We thank our program partners, including the U.S. Fish and Wildlife Service, California Department of Fish and Wildlife, the U.S. Forest Service, the Los Angeles Zoo, Omaha’s Henry Doorly Zoo and Aquarium, and the U.S. Geological Survey (USGS). We are especially grateful to Michelle Curtis for assistance with data collection and Dr. Ronald Swaisgood for his support and contribution to the broader Mountain yellow-legged Frog conservation program, which provided critical context for this work. Finally, we respectfully acknowledge the Cahuilla, Serrano, and Kumeyaay (Diegueño) peoples as the traditional custodians of the lands where portions of this research were conducted. The Cahuilla are the traditional stewards of the San Jacinto and San Bernardino Mountains, the Coachella Valley, and adjacent desert regions. The Serrano have longstanding connections to the San Bernardino and San Gabriel Mountains and the surrounding Mojave Desert margins. The Kumeyaay are the traditional custodians of the lands encompassing present-day San Diego County and extending into northern Baja California. We honour their Elders past, present, and emerging, and acknowledge the deep and enduring cultural, spiritual, and ecological relationships that connect these communities to their ancestral lands.

## CRediT authorship contribution statement

NEC, LEJ and SC designed and implemented this experiment; RU assisted with statistical design and analysis. DMS was responsible for program oversight, including funding acquisition and administration. BSD contributed to the initial experimental design and continued advice. All authors contributed to the writing and editing of the manuscript draft; all authors gave final approval for publication.

## Use of AI Technology Disclosure

In the preparation of this manuscript, Claude AI (Claude, made by Anthropic) was utilised primarily for checking grammar and R-Script development. However, Claude AI was not involved in sourcing references or producing any of the intellectual content contained herein. The final content, analysis, and conclusions of this manuscript remain the sole intellectual property of the authors.

